# Temporally distinct roles of Aurora A in polarization of the *C. elegans* zygote

**DOI:** 10.1101/2023.10.25.563816

**Authors:** Nadia I. Manzi, Bailey N. de Jesus, Yu Shi, Daniel J. Dickinson

## Abstract

During asymmetric cell division, coordination of cell polarity and the cell cycle is critical for proper inheritance of cell fate determinants and generation of cellular diversity. In *Caenorhabditis elegans* (*C. elegans*), polarity is established in the zygote and is governed by evolutionarily conserved Partitioning defective (PAR) proteins that localize to distinct cortical domains. At the time of polarity establishment, anterior and posterior PARs segregate to opposing cortical domains that specify asymmetric cell fates. Timely establishment of these PAR domains requires a cell cycle kinase, Aurora A (AIR-1 in *C.elegans*). Aurora A depletion by RNAi causes a spectrum of phenotypes including no posterior domain, reversed polarity, and excess posterior domains. How depletion of a single kinase can cause seemingly opposite phenotypes remains obscure. Using an auxin-inducible degradation system, drug treatments, and high-resolution microscopy, we found that AIR-1 regulates polarity via distinct mechanisms at different times of the cell cycle. During meiosis I, AIR-1 acts to prevent the formation of bipolar domains, while in meiosis II, AIR-1 is necessary to recruit PAR-2 onto the membrane. Together these data clarify the origin of the multiple polarization phenotypes observed in RNAi experiments and reveal multiple roles of AIR-1 in coordinating PAR protein localization with the progression of the cell cycle.

## Introduction

Cell polarity is a fundamental property of animal cells and is essential for cell fate specification and tissue formation in developing embryos. In cells that divide asymmetrically to produce daughter cells with different fates, polarity is established by proteins that localize to distinct cortical domains and promote the segregation of cell fate determinants (Boyd et al., 1996; Gönczy, 2008; Hirate et al., 2013; Knoblich, 2001; Korotkevich et al., 2017; Leung et al., 2016). For proper asymmetric cell division, the processes of cortical polarity establishment, cell fate segregation, and progression through mitosis need to be coordinated. In several well-studied cell types, cell cycle kinases directly regulate polarity proteins to ensure the coordination of cell polarization and mitotic progression (Carvalho et al., 2015; Cowan and Hyman, 2006; Dickinson et al., 2017; Noatynska et al., 2013; Wirtz-Peitz et al., 2008). Yet despite these examples, in most cases the mechanisms that coordinate cell polarity and cell cycle timing remain unclear.

The *C.elegans* zygote is a powerful model for studying asymmetric cell division and its coordination with the cell cycle. This cell establishes anterior-posterior polarity by segregating highly conserved Partitioning defective (PAR) proteins to the opposite sides of the cell (Etemad-Moghadam et al., 1995; Guo and Kemphues, 1995; Hung and Kemphues, 1999; Kemphues et al., 1988; Tabuse et al., 1998). Before polarity is established, PAR proteins are symmetrically localized and are incompetent to polarize until the zygote has completed meiosis II (Reich et al., 2019). During oogenesis, the future posterior PARs (pPARs; PAR-1, PAR-2) occupy the membrane and the future anterior PARs (aPARs; PAR-3, PAR-6, and atypical kinase C [aPKC]) remain cytoplasmic. After fertilization, aPARs gradually accumulate at the cortex around anaphase I. aPARs displace pPARs during this stage and occupy the entire cell cortex until the embryo completes meiosis II and is ready to polarize (Figure 1A, (Lang and Munro, 2017; Motegi and Seydoux, 2013; Reich et al., 2019)

Soon after meiosis II, polarization is triggered by a cell cycle kinase, Aurora A (AIR-1 in *C.elegans*) that is enriched on sperm-derived centrosomes (Figure 1A; (Kapoor and Kotak, 2019; Klinkert et al., 2019; Reich et al., 2019; Zhao et al., 2019)). AIR-1 is a conserved serine/threonine kinase that has multiple functions during cell division including centrosome maturation, spindle assembly, and spindle alignment (Hannak et al., 2001; Magnaghi-Jaulin et al., 2019; Reboutier et al., 2013; Sumiyoshi et al., 2015). During polarity establishment, recent evidence suggests that AIR-1 may act locally through a RhoA Guanine Nucleotide Exchange Factor, ECT-2, to inhibit actomyosin contractility at the posterior end of the zygote (Gan and Motegi, 2021; Kapoor and Kotak, 2019; Klinkert et al., 2019; Longhini and Glotzer, 2022; Zhao et al., 2019). The resulting anterior-directed cortical flows then sweep membrane-bound aPARs to the anterior (Chang and Dickinson, 2022; Illukkumbura et al., 2023; Munro et al., 2004). Concomitantly, PAR-2 is recruited to the posterior membrane in a way that partially depends on microtubule binding (Motegi et al., 2011). At the posterior membrane, PAR-2 then recruits the posterior kinase PAR-1 (Ramanujam et al., 2018). Polarized domains are subsequently maintained through mutual antagonism. At the anterior, aPKC phosphorylates several posterior PARs excluding them from the anterior cortex (Hao et al., 2006; Hoege et al., 2010; Kumfer et al., 2010; Motegi et al., 2011; Sailer et al., 2015). At the posterior domain, multiple pathways act together to prevent aPARs posterior association (Beatty et al., 2010; Munro et al., 2004; Sailer et al., 2015).

Although AIR-1 has been shown to regulate the anterior-posterior polarity of the *C. elegans* zygote, the exact mechanism by which AIR-1 regulates PAR protein behaviors remains unknown. AIR-1’s role has remained unclear in part because RNA interference (RNAi)-mediated depletion of AIR-1 has resulted in multiple different polarization phenotypes in different studies (Kapoor and Kotak, 2019; Klinkert et al., 2019; Reich et al., 2019; Zhao et al., 2019). The two major phenotypes are a bipolar phenotype, where pPARs are found at both ends of the embryo, and reverse polarization, where the pPAR domain forms at the maternal rather than the paternal pole. Two studies also reported that some *air-1(RNAi)* embryos completely fail to polarize: pPARs remain cytoplasmic while aPARs are uniformly distributed on the cortex (Klinkert et al., 2019; Zhao et al., 2019). These results raise a question of why similar AIR-1 depletion can lead to seemingly opposite phenotypes: one where PAR-2 forms excess cortical domains (bipolar), and another where PAR-2 does not reach the cortex at all.

To dissect the role of AIR-1 in regulating pPAR cortical recruitment and understand the basis for these different phenotypes, we used a combination of auxin-inducible degradation, drug treatment, and high-resolution microscopy to interfere with AIR-1 function at different times of the cell cycle and observe how PAR polarity is affected. We found that AIR-1 regulates polarity via different mechanisms at different times. AIR-1 is required during meiosis I to prevent the formation of bipolarity and reversed polarization, while AIR-1 catalytic activity during meiosis II is required for PAR-2 to load on the membrane. The early function of AIR-1 in suppressing bipolarity involves another mitotic kinase, PLK-1, but the late function in symmetry breaking does not. Our work helps clarify the origin of what had been a puzzling spectrum of phenotypes resulting from AIR-1 depletion and sheds light on the role of AIR-1 in establishing the anterior-posterior polarity of a *C.elegans* zygote.

## Results

### Depleting Aurora A using RNAi or kinase inhibitor shows distinct phenotypes

Previous studies reported multiple phenotypes, in different proportions, following *air-1* RNAi treatment. (Kapoor and Kotak, 2019; Klinkert et al., 2019) reported the formation of mostly bipolar embryos (PAR-2 formed both anterior and posterior domains), whereas (Reich et al., 2019; Zhao et al., 2019) observed similar amounts of reverse (PAR-2 formed a domain at the anterior) and bipolar phenotypes. Additionally, some studies reported a complete loss of PAR-2 domains in a minority of embryos following *air-1* RNAi (Klinkert et al., 2019; Zhao et al., 2019). We therefore began by repeating the previous experiment: we depleted AIR-1 using RNAi and performed confocal live imaging in a strain carrying endogenously tagged mNeonGreen(mNG)::PAR-2 and PAR-6::mScarlet(mSc) (Figure 1B). In our hands, 54% of *air-1(RNAi)* embryos were bipolar, 11% were reversed, 29% were normally polarized, and 7% failed to form a PAR-2 domain (Figure 1B-C).

To understand why depleting AIR-1 leads to these distinct PAR-2 behaviors, we first sought to test the hypothesis that variation in phenotypes may result from differences in RNAi penetrance. To test whether there was residual AIR-1 after RNAi, we endogenously tagged AIR-1 with mNG for visualization and fluorescent intensity measurements. Indeed, we found that *air-1(RNAi)* embryos contained residual mNG::AIR-1, which was visible as a faint signal on centrosomes at the onset of cytokinesis (Figure 1D). If variation in the amount of residual AIR-1 could account for the variable phenotypes, then embryos with different polarity phenotypes should have different amounts of residual AIR-1. However, we measured AIR-1 fluorescent intensities around centrosomes and found that all *air-1(RNAi)* embryos had similar amounts of residual AIR-1, regardless of their polarity phenotype (Figure 1E). This result is not consistent with the hypothesis that different *air-1(RNAi)* phenotypes result from different levels of AIR-1 depletion.

We next tested AIR-1 inhibition using chemical inhibitors, because we reasoned that titrating drug concentration might allow us to better control the amount of residual AIR-1 activity and explore whether this could account for phenotypic variability. Because Aurora A is highly conserved among species (Brown et al., 2004), we used commercially available Aurora A kinase inhibitors (Sells et al., 2015) to perturb AIR-1 kinase activity. We permeabilized embryos’ eggshells using RNAi against the eggshell component *perm-1* (Carvalho et al., 2011; Olson et al., 2012). Permeabilized embryos were dissected into Shelton’s growth media (SGM) containing either dimethyl sulfoxide (DMSO) or an Aurora A kinase inhibitor (MLN8054) and immediately mounted for live imaging. 100% of control DMSO-treated embryos established a single polarized PAR-2 domain by pronuclear meeting (PNM) (Figure 1F). Surprisingly, the majority of embryos treated with 20μM MLN8054 failed to localize PAR-2 on the membrane during polarity establishment and never polarized throughout the maintenance phase (Figure 1G-H). We saw only a single bipolar embryo among those treated with 20μM MLN8054, and the anterior PAR-2 domain in this embryo was very small and co-localized with PAR-6 (Figure S1A). We referred to this phenotype as “weak bipolar” since it is distinct from the bipolar phenotype observed in *air-1(RNAi)* (Figure 1B, Figure S1B). Thus, unexpectedly, AIR-1 inhibition causes a different phenotype than AIR-1 depletion via RNAi.

**Figure 1.**
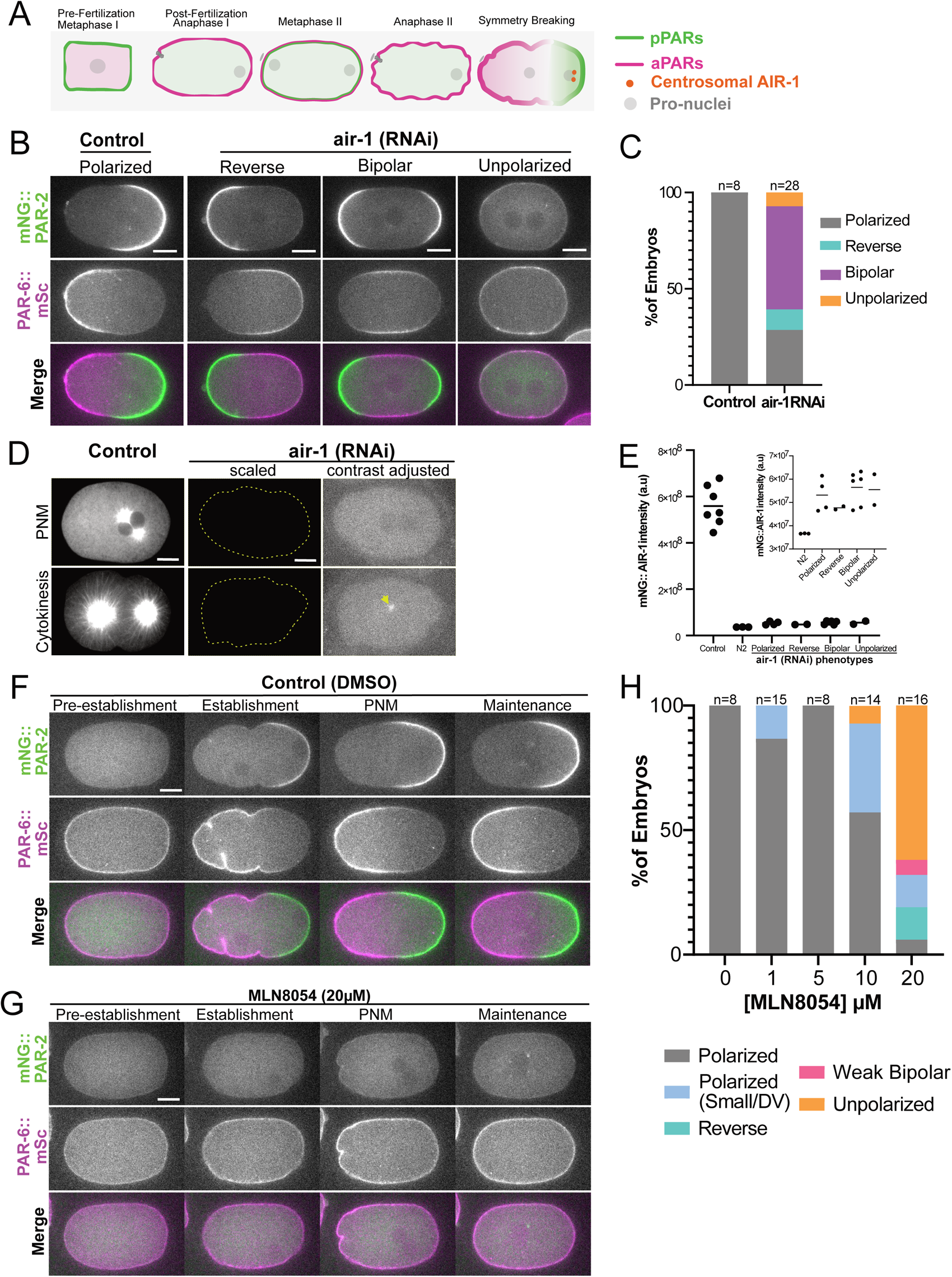
Inhibition of AIR-1 kinase activity shows distinct phenotypes from RNAi-mediated depletion. **A:** Schematic showing PAR proteins localization from oocyte to zygote transition. In oocytes posterior PARs (pPARs) localize to the membrane while anterior PARs (aPARs) are in the cytoplasm. By anaphase I, aPARs load and remain on cortex until the embryo completes meiosis II. pPARs become restricted in the cytoplasm after fertilization, briefly load on the membrane at metaphase I but re-localize to the cytoplasm until the embryo is ready to polarize. After meiosis II, centrosomal AIR-1 triggers symmetry breaking: aPARs segregate to the anterior, pPARs load to the posterior domain. **B**: Mid-plane images from time-lapse live imaging of mNG::PAR-2; PAR-6::mSc embryos in control or air-1 RNAi conditions showing multiple polarization phenotypes. Scale bar represents 10μm **C**: Quantification of polarization phenotypes from panels in B scored at pronuclear meeting (PNM). **D**: Mid-plane images from time-lapse live imaging of mNG::AIR-1 embryos in either control (left panel) or *air-1 RNAi* conditions (middle and right panel). The middle panel’s contrast is scaled to match the control (the outline of the embryo is in yellow). The right panel’s contrast is adjusted to increase visibility. Scale bar represents 10μm **E**: Quantification of AIR-1 fluorescence intensity from data in D. A rectangular box was drawn around the centrosomes and the total pixel intensity minus cytoplasmic background was measured in FIJI. Inset represents mNG::AIR-1 fluorescence intensity in embryos exhibiting different phenotypes compared to background fluorescence in a strain with no tag (N2). **F-G**: Montage from time-lapse live imaging of permeabilized mNG::PAR-2; PAR-6::mSc embryos treated with 0.2% DMSO control or 20 μM of Aurora A inhibitor, MLN8054. Scale bar represents 10μm **H**: Quantification of phenotypes in panels F-G with additional examples of embryos treated with different MLN8054 concentrations. Phenotypes were scored at PNM

We next titrated the amount of Aurora A inhibitor to determine whether embryos with partial inhibition of AIR-1 kinase activity would phenocopy *air-1(RNAi)* embryos. However, even at lower concentrations, we did not see embryos with bipolar PAR-2 domains (Figure 1H). Instead, some embryos polarized but formed small posterior or dorsal/ventral (D/V)-localized PAR-2 domains (Figure S1A, Figure 1H). To confirm these results, we used a different Aurora A kinase inhibitor, alisertib (MLN8237). 86% of embryos treated with 20μM MLN8237 failed to polarize, and only 14% were bipolar (Figure S1D-E). At lower MLN8237 concentrations, we did observe a minority of embryos that formed weak bipolar domains. In these weak bipolar embryos, PAR-2 was absent from the membrane during polarity establishment (i.e., prior to PNM) but formed small bipolar domains during maintenance phase that overlapped with PAR-6 (Figure S1E-F).

To exclude the possibility that the failure of PAR-2 to bind to the membrane is a result of off-target effects of the Aurora A inhibitors, we performed RNAi against AIR-1’s closest paralog, Aurora B / AIR-2. Embryos depleted of AIR-2 polarized normally and embryos co-depleted of AIR-1 and AIR-2 formed 79% polarized domains, 17% bipolar, and only 4% unpolarized phenotypes (Figure S1G). The milder phenotype resulting from AIR-1 and AIR-2 co-depletion compared to *air-1(RNAi)* alone is likely a result of reduced RNAi penetrance when targeting multiple genes, a well-known issue with RNAi experiments in *C. elegans.* Although we cannot formally rule out off-target effects of Aurora A inhibitors on other kinases, these results suggest that the failure of PAR-2 to load onto the membrane is not a consequence of unintended Aurora B / AIR-2 inhibition. Altogether, these data indicated that AIR-1 catalytic activity is required to localize PAR-2 on the membrane in a timely manner and that AIR-1 inhibition and RNAi-mediated depletion produce distinct phenotypes.

### AIR-1 regulates PAR protein localization via different mechanisms at different times in development

We sought to understand why PAR-2 is recruited to the membrane (in either a polarized or bipolarized fashion) in most *air-1(RNAi)* embryos, but not in embryos where AIR-1’s kinase activity is inhibited. We first hypothesized that AIR-1 may have a non-catalytic function that regulates polarity establishment, similar to the non-catalytic roles of Aurora A in regulating meiotic and mitotic processes in different systems (Guarino Almeida et al., 2020; Toya et al., 2011). Reducing AIR-1 protein levels via RNAi would interfere with both catalytic and non-catalytic functions of AIR-1, while inhibitor treatment would block catalytic activity but leave putative non-catalytic activities intact. In this model, a non-catalytic function of AIR-1 would be required to suppress bipolarity, since bipolar embryos are observed following AIR-1 RNAi but not after inhibitor treatment. To test this idea, we imaged embryos depleted of endogenous AIR-1 via RNAi but expressing RNAi-resistant, kinase-dead GFP::AIR-1 (Klinkert et al., 2019). If a non-catalytic activity of AIR-1 suppressed bipolarity, then kinase-dead AIR-1 should retain this activity and the resulting embryos should lack polarized PAR-2 domains, similar to inhibitor-treated embryos. However, we found that embryos expressing kinase-dead AIR-1 showed phenotypes similar to those observed in *air-1(RNAi)* embryos (73% bipolar, 13% polarized, and 13% unpolarized) (Figure S1H, Figure S1I). The finding that most embryos expressing kinase-dead AIR-1 are bipolar confirms a previous report (Klinkert et al., 2019). Thus, the different phenotypes of *air-1(RNAi)* and inhibitor-treated embryos cannot be explained by non-catalytic functions of AIR-1.

We next considered the timing of AIR-1 activity. We reasoned that the depletion of AIR-1 with RNAi is a chronic treatment: dsRNA is injected into the mother and embryos are imaged 24 hours later, so the resulting phenotypes reflect a loss of AIR-1 throughout oogenesis. In contrast, kinase inhibitors are added to fertilized embryos that are exiting meiosis I or are in meiosis II, producing an acute effect. AIR-1 has been shown to have roles during oogenesis (Reich et al., 2019) and to have dynamic localization throughout the course of the zygote cell cycle (Klinkert et al., 2019), supporting the possibility that AIR-1 could regulate PAR protein localization differently at different times.

To test this hypothesis, we used an auxin-inducible degradation system (Martinez and Matus, 2020; Yesbolatova et al., 2020; Zhang et al., 2015) to degrade AIR-1 at different times of the cell cycle. In brief, we constructed a strain expressing an E3 ligase Tir1 and in which AIR-1 is endogenously tagged with an auxin-inducible degron (AID) fused to mNG. In the presence of auxin, mNG::AID::AIR-1 is polyubiquitinated and degraded via the proteasome (Figure 2A). In the absence of auxin, mNG::AID::AIR-1 was expressed and localized normally, and in most embryos, endogenously tagged mSc::PAR-2 formed a single polarized posterior domain with normal timing (Figure 2B). The bipolar embryos observed in the absence of auxin do not reflect a loss of function caused by the AID tag because we also observed occasional bipolar phenotypes in wild-type strains co-expressing endogenously tagged mNG::AIR-1 and mSc::PAR-2 (Figure S2A). We suspect that these bipolar phenotypes may result from steric collisions between the tags, since strains carrying either mSc::PAR-2 or mNG::AIR-1 alone polarize normally. To degrade mNG::AID::AIR-1, we first incubated worms with auxin for 24 hours, mimicking a typical RNAi experiment. mNG::AID::AIR-1 was reduced to undetectable levels following this treatment (Figure 2C). In a majority of the resulting embryos, PAR-2 was able to load on the membrane and formed bipolar domains (Figure 2D-E), similar to *air-1(RNAi)* embryos (Figure 1B, Figure S1B). However, unlike in RNAi experiments, we did not observe any unpolarized embryos that failed to localize PAR-2 to the membrane following 24h auxin treatment. Next, to mimic acute treatment with Aurora A inhibitors, we soaked worms in buffer containing 4mM auxin for 15 minutes, then dissected and imaged the resulting embryos. Strikingly, the effects of a 15-minute auxin treatment matched those from Aurora A inhibitor treatment: 83% of embryos failed to break symmetry and retained PAR-2 in the cytoplasm (Figure 2E-F, compare to Figure 1G). In these experiments, some mNG::AID::AIR-1 expression was visible at the beginning of the imaging, but it was gradually depleted thereafter and was reduced to background levels by PNM (Figure 2C & 2F).

**Figure 2:**
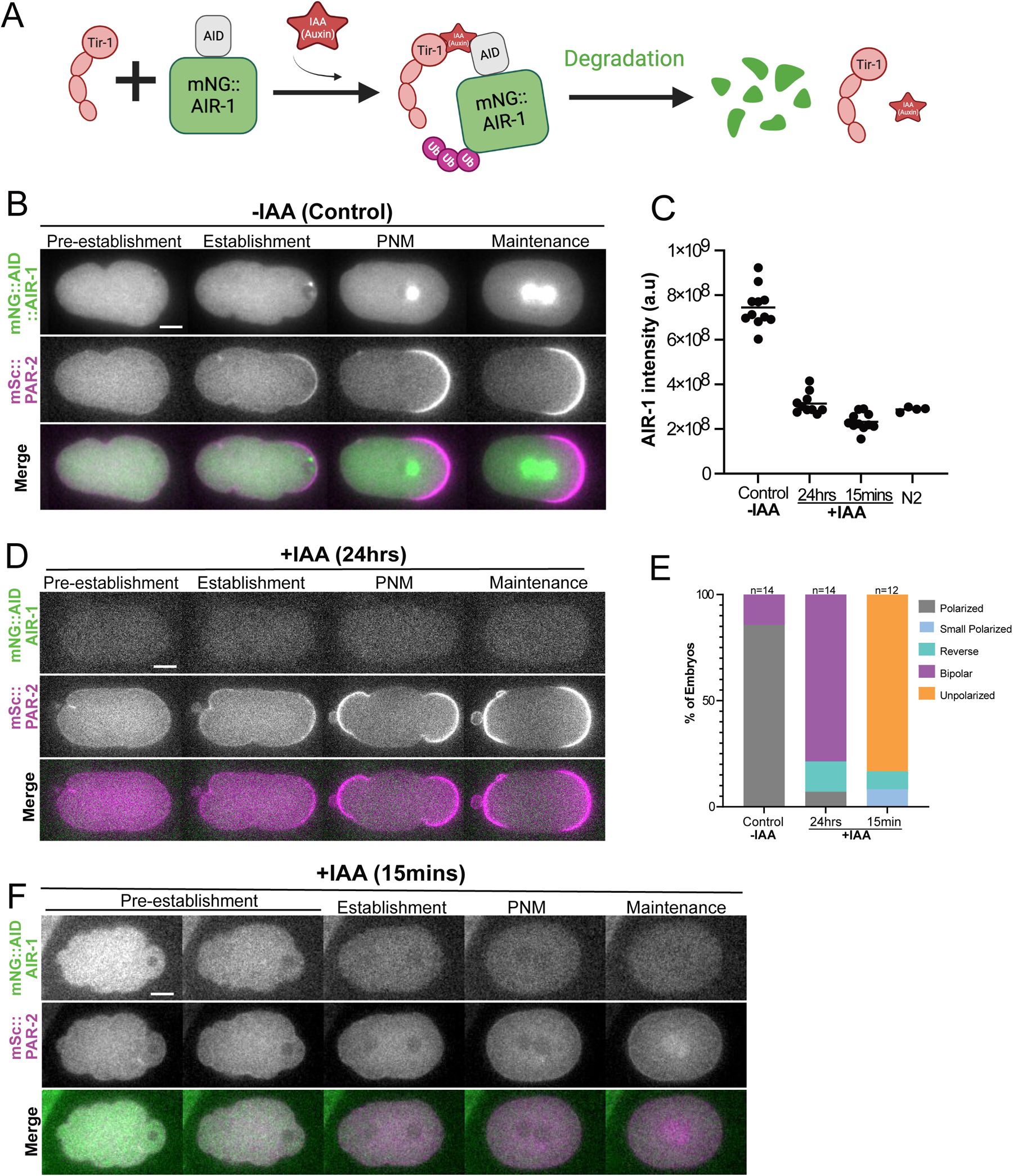
AIR-1 has distinct regulation of PAR-2 localization at different times of the cell cycle. **A**: Illustration of the Auxin-inducible degradation system. AIR-1 is endogenously tagged with a degron, AID, and mNG in a strain expressing an E3 ligase, TIR-1. In the presence of Auxin, mNG::AID::AIR-1 is polyubiquitinated and degraded via the proteasome. **B**: Montage from time-lapse live imaging of mNG::AID::AIR-1; mSc::PAR-2 embryos treated with 1% ethanol control. Scale bar represents 10μm **C**: Quantification of total AIR-1 fluorescent intensity at pronuclear meeting (PNM) from images in B, D, F, and N2 (non-fluorescently tagged embryos). **D:** Montage from time-lapse live imaging of mNG::AID::AIR-1; mSc::PAR-2 embryos treated with 4mM auxin for 24 hours. Scale bar represents 10μm **E**: Quantification of phenotypes scored at PNM from panels B, D, and F. **F**: Montage from time-lapse live imaging of mNG::AID::AIR-1; mSc::PAR-2 embryos treated with 4mM auxin for 15 minutes. Scale bar represents 10μm

We also examined aPKC localization in worms treated with auxin for 15 minutes or 24 hours. In the absence of auxin, aPKC formed a single anterior domain and AIR-1 was concentrated on the centrosomes (Figure S2B-C). In worms treated with auxin for 15 minutes, AIR-1 was degraded and aPKC remained uniformly localized on the membrane (Figure S2B-C). In contrast, embryos treated with auxin for 24 hours formed a band around the center of the embryo, indicating the presence of bipolar pPAR domains at both ends (Figure S2B-C).

Together, these data suggest distinct functions for AIR-1 at different stages of development. AIR-1 acts early in development, most likely in the maternal germline prior to meiosis, to suppress the formation of bipolar PAR-2 domains. After fertilization, AIR-1 is necessary to allow PAR-2 to load on the membrane and establish a single polarized domain.

To better define the functions of AIR-1 at different stages of oogenesis and the cell cycle, we depleted AIR-1 via the AID system while performing *in utero* live imaging. This approach allowed us to deplete AIR-1 at different times prior to symmetry breaking and test how polarity establishment was affected. We first imaged mNG::AID::AIR-1; mSc::PAR-2 embryos in the absence of auxin. We followed mSc::PAR-2 localization over time and used the fact that mNG::AID::AIR-1 localizes at meiotic spindles and chromosomes of a newly fertilized embryo to determine the exact stage of meiosis. Consistent with a previous report (Reich et al., 2019), PAR-2 was localized uniformly on the membrane immediately after fertilization (-36mins relative to PNM) but was cleared by anaphase I. At metaphase II (-19:30 mins prior to PNM), PAR-2 was transiently bound to the membrane, but it relocalized to the cytoplasm prior to symmetry breaking. Soon after the end of meiosis II, 9 of 10 embryos formed a single polarized posterior PAR-2 domain (Figure 3A) and 1/10 embryos formed bipolar PAR-2 domains (not shown).

In embryos treated with auxin before or during meiosis I (-30mins or earlier relative to PNM), PAR-2 localized either in a bipolar fashion (Figure 3Bi, Figure 3C, Figure S3A) or at the maternal (normally anterior) membrane (Figure 3Bii, Figure 3C, Figure S3B). The frequency of reverse polarity was increased *in utero* compared to *ex utero* embryos from worms that were treated with auxin or RNAi for 24 hours (Figure 3C, compared to Figure 1C, Figure 2D-E), in agreement with a previous study (Reich et al., 2019). We speculate that this difference is due to the different embryo environments (see Discussion). In more than half of embryos depleted of AIR-1 in meiosis II (-30mins or later relative to PNM), PAR-2 failed to load on the posterior membrane by PNM (Figure 3Biii, Figure 3C, Figure S3C). These results are similar to the phenotype observed when post-fertilization embryos are treated with Aurora A kinase inhibitors (Figure 1G-H).

We estimated the time of AIR-1 depletion relative to PNM and observed that loss of AIR-1 during meiosis I, which starts during oogenesis and ends quickly after fertilization, results in the bipolar or reverse polarization of PAR-2, whereas loss of AIR-1 during meiosis II mostly leads to failure of PAR-2 to load on the membrane (Figure 3C).

**Figure 3:**
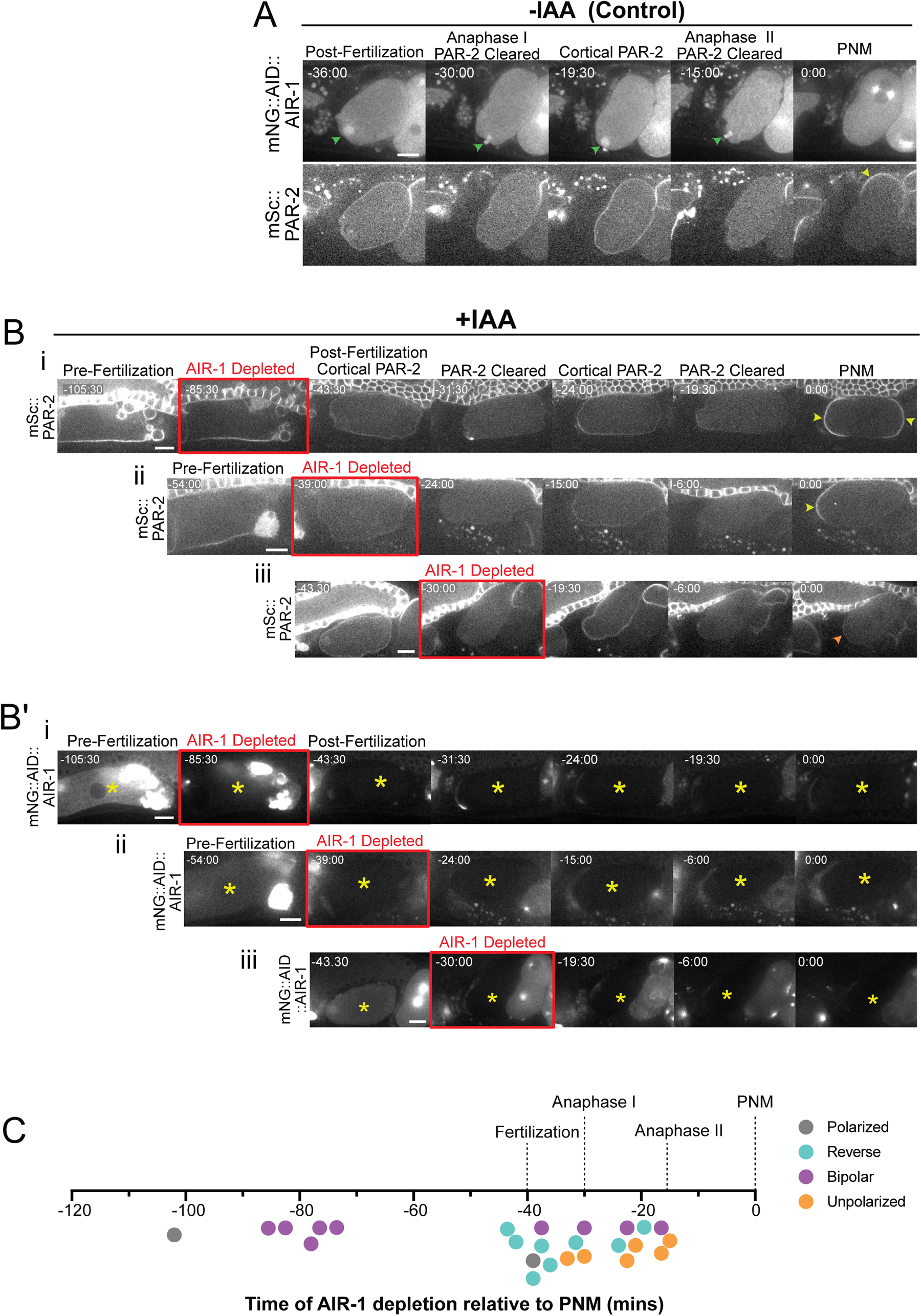
AIR-1 in meiosis II not meiosis I is required for PAR-2 membrane localization and symmetry breaking. **A**: Montage from time-lapse live imaging of mNG::AID::AIR-1; mSc::PAR-2 embryos. Embryos were imaged *in utero* soon after fertilization from worms that were mounted in M9 without Auxin (IAA). Green arrowhead indicate meiotic DNA and spindle organization used to estimate meiosis stage, while yellow arrowhead shows PAR-2 membrane localization at the time of pronuclear meeting (PNM). The time scale shown is in minutes and relative to PNM. Scale bar represents 10μm **B-B’**. Montage from time-lapse *in-utero* live imaging of worms expressing mSc::PAR-2 (**B**) and mNG::AID::AIR-1 (**B’**). Embryos were imaged starting from before fertilization (**i** and **ii**) or after fertilization (**iii**) from worms that were mounted in 4mM auxin (IAA) buffer. The time of AIR-1 depletion is shown in red. The resulting phenotypes after AIR-1 depletion show bipolar (**i**), reverse (**ii**) and unpolarized (**iii**). Yellow arrowheads indicate PAR-2 membrane localization while orange arrowheads indicate embryos lacking cortical PAR-2. Yellow asterisks indicate the representative embryo. The time scale is shown in minutes relative to pronuclear meeting. Scale bar represents 10μm **C**: Plot showing the time (in minutes) by which AIR-1 was depleted relative to pronuclear meeting and the resulting polarization phenotype. Each data point represents one embryo observed *in utero.* The position on the timeline indicates the time when AIR-1 was effectively depleted, and the color of the point indicates the observed phenotype.

Together these data reveal two distinct roles of AIR-1 in regulating *C. elegans* zygote polarity. First, during meiosis I, AIR-1 inhibits premature PAR-2 loading onto the anterior cortex, which leads to reversed or bipolar phenotypes. Second, AIR-1 plays a positive role that is required for symmetry breaking and PAR-2 membrane localization in the zygote following the completion of meiosis II.

### PLK-1 contributes to restraining polarity during meiosis I but is dispensable for symmetry breaking

Our data up to this point suggest two distinct functions for Aurora A in regulating polarity: An early function that suppresses reverse and bipolar phenotypes, and a late function that is required for PAR-2 loading on the cortex. Previously, the activity of AIR-1 that suppresses reverse and bipolarity has been proposed to be shared with another mitotic kinase, Polo-like kinase, PLK-1, (Noatynska et al., 2010; Reich et al., 2019). Depletion of either AIR-1 or PLK-1 via RNAi produces premature symmetry breaking and a mixture of reverse and bipolar phenotypes. We therefore wondered whether PLK-1, too, has a later function that directly promotes PAR-2 cortical loading and symmetry breaking.

To look for a role of PLK-1 in symmetry breaking, we used a specific inhibitor, BI2536, to acutely block PLK-1 activity (Steegmaier et al., 2007) in fertilized embryos entering meiosis II and tested whether symmetry breaking was affected. First, to confirm that this inhibitor blocks PLK-1 activity in *C. elegans* embryos, we imaged embryos treated with BI2536 expressing mNG::PAR-3, an aPAR whose clustering on the cell cortex is negatively regulated by PLK-1 (Chang and Dickinson, 2022; Dickinson et al., 2017). Control embryos dissected in DMSO formed PAR-3 oligomers that segregated to the anterior domain. These clusters dissolved during polarity maintenance, and PAR-3 membrane localization was reduced (Figures 4A & 4C). Embryos treated with 20 μM BI2536 formed anterior PAR-3 oligomers that persisted throughout maintenance phase (Figures 4B $ 4C), a phenotype similar to that observed in *plk-1 (RNAi)* embryos (Dickinson et al., 2017). To determine how PAR-2 localization was affected in the absence of PLK-1 kinase activity, we repeated the same experiment in a strain expressing mNG::PAR-2. Interestingly, we found that in embryos treated with PLK-1 inhibitors, PAR-2 was recruited to the membrane and formed either polarized PAR-2 domains, with the posterior domain sometimes expanded abnormally to the anterior (Figure 4D-group1, Figure 4E), or transient weak bipolar phenotypes, where PAR-2 formed a small anterior domain that cleared by PNM (Figure 4D-group2, Figure 4E). These phenotypes were similar to phenotypes reported in *plk-1 RNAi* embryos (Calvi et al., 2022; Noatynska et al., 2010). These results indicate that PLK-1 catalytic activity is dispensable in recruiting PAR-2 to the posterior membrane.

**Figure 4:**
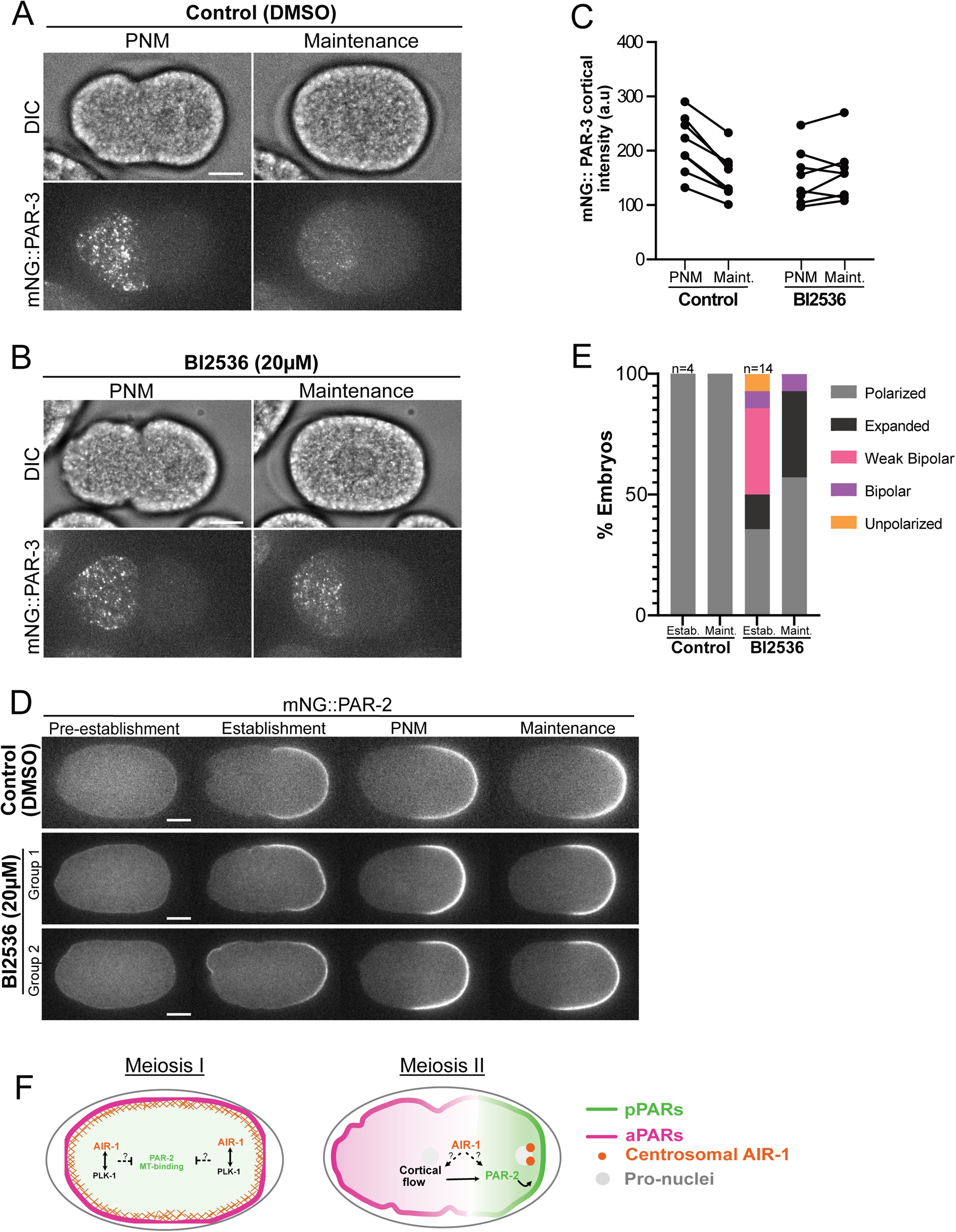
PAR-2 Cortical recruitment is independent of PLK-1 kinase activity. **A-B** : Cortical maximum intensity projection images from time-lapse live imaging of permeabilized embryos expressing mNG::PAR-3 treated with 0.01% DMSO (**A**) or PLK-1 kinase inhibitor, BI2536 (**B**) at pronuclear meeting (Left) or maintenance (right). DIC images are shown in the top panels. Scale bar represents 10μm **C**. Quantification of mNG::PAR-3 cortical fluorescent intensity from data in A-B at pronuclei meeting (PNM) or Maintenance (5 mins post-PNM). Mean cortical intensities were measured in FIJI by drawing a box in the anterior (left) side of the embryo and subtracting off-embryo background. **D**: Montage from time-lapse live imaging of permeabilized embryos expressing mNG::PAR-2 treated with 0.01% DMSO or 20 μm of PLK-1 kinase inhibitor, BI2536. The phenotypes of embryos lacking PLK-1 kinase activities were categorized in two main groups. In group 1 (middle panel) embryos, PAR-2 forms a domain at the posterior that extends to the anterior at polarity establishment but this domain is corrected by pronuclear meeting. In group 2 (bottom panel) embryos, PAR-2 forms bipolar domains at establishment. The anterior domain clears and the posterior domain extends to the anterior compared to the control (top panel). Scale bar represents 10μm **E**: Quantification of PAR-2 phenotypes of data from D. **F**. Schematic showing the current proposed working model. In meiosis I, cortical AIR-1 works in concert with PLK-1 to inhibit PAR-2 microtubule association, which in-turn may prevent premature formation of bipolarization or reverse polarity. In meiosis II, centrosomal AIR-1 acts directly through PAR-2 or actomyosin cortical flow to promote recruitment of PAR-2 at the anterior membrane thus allowing formation of a single polarized domain.

We conclude that although AIR-1 and PLK-1 may cooperate to suppress formation of ectopic polarized domains (Kapoor and Kotak, 2019; Klinkert et al., 2019; Reich et al., 2019; Zhao et al., 2019), AIR-1 acts independently of PLK-1 to initiate symmetry breaking in the zygote.

## Discussion

Coupling of cell polarity with the cell cycle is critical for the development of multicellular organisms. Previous work showed that anterior-posterior polarity in the *C. elegans* zygote is initiated by Aurora A (AIR-1 in *C. elegans*) (Kapoor and Kotak, 2019; Klinkert et al., 2019; Reich et al., 2019; Zhao et al., 2019). However, depletion of AIR-1 by RNAi caused a spectrum of distinct phenotypes including bipolarity (PAR-2 forms an anterior and a posterior domain), reverse polarity (PAR-2 localizes in the anterior cortex), and no polarization (PAR-2 fails to load to the membrane), raising the question of how AIR-1 regulated PAR-2 localization and/or cortical recruitment. Here, we have presented evidence that AIR-1 regulates PAR polarity via different mechanisms at different times of the cell cycle. In meiosis I, AIR-1 prevents premature cortical localization of PAR proteins (Reich et al., 2019) and thereby contributes to the timely and precise formation of single polarized domains. In meiosis II, AIR-1 is required to load PAR-2 on the membrane in a manner that depends on its kinase activity and is independent of PLK-1.

Mechanistically, how does AIR-1 regulate PAR-2 localization? PAR-2 contains several Aurora A consensus phosphorylation motifs, comprising arginine/lysine residues at the -2 or -3 position and leucines in the -1 and +1 positions relative to the targeted serine (Hao et al., 2006; Kettenbach et al., 2011; Ramanujam et al., 2018). Thus, a simple hypothesis is that AIR-1 might directly phosphorylate PAR-2, which could either promote or inhibit its recruitment into cortical domains (Hao et al., 2006). This hypothesis is not straightforward to test *in vivo* because AIR-1 and aPKC have similar phosphorylation motifs (Kettenbach et al., 2011; Kreegipuu et al., 1998; Nishikawa et al., 1997; Ramanujam et al., 2018) and might even phosphorylate some of the same sites. Phosphorylation of PAR-2 by aPKC is already known to inhibit its cortical localization, suggesting that AIR-1 might play a similar role, but careful biochemical experiments will be required to test this in detail.

Consistent with the possibility that PAR-2 may be a direct AIR-1 substrate, we have observed that some knock-in strains carrying fluorescent tags on both AIR-1 and PAR-2 – but not AIR-1 in combination with other PARs – form bipolar PAR-2 domains *in utero* and *ex utero,* similar to embryos depleted of AIR-1 with RNAi or 24h auxin treatment (Figure S2A). Since single-labeled AIR-1 or PAR-2 knock-in strains do not exhibit a bipolar phenotype, we speculate that it might result from steric hindrance that prevents tagged AIR-1 and tagged PAR-2 from interacting normally.

Alternatively or in addition, AIR-1 may modulate PAR-2 microtubule binding. PAR-2 localizes to the membrane through two semi-redundant pathways: cortical flows that clear aPARs from the posterior domain (Chang and Dickinson, 2022; Cheeks et al., 2004; Goehring et al., 2011; Illukkumbura et al., 2023; Munro et al., 2004) allowing PAR-2 to associate with the posterior domain, and a second pathway involving microtubule binding which protects PAR-2 from aPKC phosphorylation (Motegi and Seydoux, 2013; Motegi et al., 2011). The microtubule-dependent pathway is required for bipolar PAR-2 domain formation in the absence of AIR-1 (Klinkert et al., 2019) suggesting a model in which AIR-1 acts during meiosis I to inhibit PAR-2 from binding microtubules. We note that although a previous study reported that nocodazole-treated *air-1(RNAi)* embryos still formed bipolar PAR-2 domains (Kapoor and Kotak, 2019), these results do not rule out a microtubule-dependent mechanism for AIR-1 in polarity because nocodazole was added only at meiosis II, which is after the stage when we show AIR-1 is acting to suppress bipolarity. Later, after meiosis II, AIR-1 most likely causes symmetry breaking through a microtubule-independent route (Figure 4F).

Another possibility is that AIR-1 regulates PAR-2 membrane localization via indirect mechanisms. In other systems, anterior PAR proteins have been shown to be Aurora A substrates: for example, in drosophila neuroblasts, Aurora A phosphorylates PAR-6 to allow numb basal localization (Wirtz-Peitz et al., 2008). In mammalian cells, Aurora A phosphorylates PAR-3 to promote neuronal polarity (Khazaei and Püschel, 2009). Since PAR-3, PAR-6, and aPKC form the aPAR complex that is antagonistic to pPAR cortical localization, AIR-1 could promote PAR-2 localization by antagonizing aPARs. Looking outside the PAR system, a recent study showed that PAR-2 membrane localization requires PP1 phosphatases GSP-1 and GSP-2 (Calvi et al., 2022). Centrosomal AIR-1 could promote GSP1/2 activation by interacting with and/or phosphorylating GSP1/2 itself or its regulatory binding partners.

Depletion of AIR-1 for 24 hours by RNAi (Kapoor and Kotak, 2019; Klinkert et al., 2019; Reich et al., 2019; Zhao et al., 2019) or auxin-inducible degradation system (this study) leads to multiple polarization phenotypes. Although the frequency of these phenotypes varied among the different studies and different strains, prominent phenotypes observed in *ex utero* embryos were characterized by the formation of more bipolar phenotypes compared to reverse polarity (Figure 1, Figure 2). However, we observed that embryos depleted of AIR-1 *in utero* showed less bipolarity and instead shifted toward reverse polarization (Figure 3, (Reich et al., 2019; Zhao et al., 2019)). One possible explanation is that *in utero* embryos depleted of AIR-1 exist in their native environment and are exposed to extracellular signals that suppress bipolarity via a different mechanism. Another possibility is that bipolarity or reverse polarization observed when mNG::AID::AIR-1 is degraded during meiosis I is a secondary effect of failed meiotic division. AIR-1 plays an important role in meiotic progression in *C. elegans* and mammals (Blengini et al., 2021; Saskova et al., 2008; Sumiyoshi et al., 2015) and embryos arrested in meiosis I segregate PAR-2 to the anterior (Reich et al., 2019; Wallenfang and Seydoux, 2000).

We have also shown that during symmetry breaking, membrane localization of PAR-2 does not require polo-like kinase (PLK-1) activity. Previous work has suggested a model in which AIR-1 acts through PLK-1 to promote the formation of a single posterior PAR-2 domain (Reich et al., 2019). However, we found that in embryos treated with PLK-1 inhibitors, PAR-2 retained its ability to bind the membrane and showed weak bipolar and extended domains phenotypes, similar to those observed in *plk-1(RNAi)* (Calvi et al., 2022; Noatynska et al., 2010; Reich et al., 2019). Because AIR-1 has multiple functions and substrates during the different stages of the cell cycle (Kettenbach et al., 2011; Magnaghi-Jaulin et al., 2019; Sardon et al., 2010; Tien et al., 2004), it is possible that AIR-1 works in concert with PLK-1 early in meiosis I to prevent microtubule-binding dependent PAR-2 cortical recruitment whereas, in meiosis II, AIR-1 acts through a novel target that is yet to be identified (Figure 4F).

In summary, our work has demonstrated temporally distinct roles of Aurora A kinase in regulating *C. elegans* anterior-posterior polarization. During meiosis I, AIR-1 ensures the formation of single posterior PAR-2 domains. In meiosis II, AIR-1 is required to recruit PAR-2 on the membrane through mechanisms that are yet to be investigated. Aurora A is conserved among different species and plays critical roles in cell division and cell polarity. Deciphering the mechanisms by which Aurora A regulates polarity proteins at different times of the cell cycle may contribute to the understanding of how polarity is coordinated with the cell cycle in other asymmetrically dividing cells.

## Author Contributions

NIM and DJD conceived of the project and designed the experiments. NIM performed and analyzed experiments in Figure 1 and Figure 3, BND performed and analyzed experiments in Figure 4, NIM and BND performed and analyzed experiments in Figure 2. YS constructed the mSc::PAR-2 allele. DJD supervised the project and secured funding. NIM and DJD co-wrote the manuscript, and all authors discussed and contributed to the final version.

## Acknowledgments

We thank Pierre Gönczy for sharing *C. elegans* strains, Monica Gotta for helpful discussions, and all the members of the Dickinson Lab for their critical feedback on the manuscript. This work was supported by the U.S. National Institutes of Health (R01 GM138443) and by a grant from the Mallinckrodt Foundation. NIM was supported by an NIH predoctoral fellowship (F31 HD108006). DJD is a CPRIT Scholar supported by the Cancer Prevention and Research Institute of Texas (RR170054).

## Materials and Methods

### Materials, organisms, and software

Reagents, strains, and software are listed in the key resource table

### *C. elegans* strain construction and maintenance

All strains were fed *E-coli* OP50 and maintained on nematode growth medium (NGM) plates at 20℃.

Genetic modification including fluorescent protein knock-ins with/without degron AID was performed using CRISPR/Cas9-triggered homologous recombination following protocols published by our laboratory (Dickinson et al., 2013; Dickinson et al., 2015; Huang et al., 2021)

### Methods details

#### Sample preparation

*Ex-utero* embryos were dissected from gravid adults in 10 µL of 1X egg buffer (5 mM HEPES pH 7.4, 118 mM NaCl, 40 mM KCl, 3.4 mM MgCl2, 3.4 mM CaCl2) containing 22.8 µm beads (Whitehouse Scientific, Chester, UK) to act as spacers, mounted on a poly-lysine coated 22×22 µm glass coverslip and sealed with VALAP (1:1:1 vaseline:lanolin:paraffin wax)

*In-utero* embryos were imaged from whole worms soaked in 0.1 μm polystyrene beads (Polysciences) diluted M9 buffer (22.5mM KH_2_PO_4_, 42.5mM Na_2_HPO_4_, 86mM NaCl, 1mM MgSO_4_) at 1% concentration, mounted with 22×22 µm glass coverslip and agar pad (7.5% agarose in M9) and sealed with VALAP (1:1:1 vaseline:lanolin:paraffin wax)

#### Confocal microscopy

Images in Figure 3 and Figure S3 were taken on a Nikon Ti2 microscope controlled by Micro-Manager software and equipped with a 60x, 1.4 NA oil immersion objective lens; an X-Light V3 spinning disk confocal head (Crest Optics, Rome, Italy); and a Prime95B sCMOS camera (Teledyne Photometrics, Tucson, AZ). All other images were taken by a Nikon Ti2 microscope controlled by Micro-Manager software and equipped with a 60x, 1.4 NA oil immersion objective lens; OptoSpin filter wheel (CAIRN Research, Kent, England), an iSIM super-resolution confocal scan head (Visitech, Sunderland, UK); and a Teledyne photometrics PrimeBSI or Kinetix22 camera. GFP and mNG were excited by 488 nm or 505 nm lasers, while mScarlet-I and mCherry were excited by 555 nm or 561 nm lasers, and appropriate single-bandpass emission filters were used for detection.

#### RNA interference

RNA interference (RNAi) was performed by injection (Figure 1I) or by feeding (all other experiments).

To perform RNAi by injection, we amplified 0.5-2kb of the target gene from N2 cDNA using primers containing T7 promoters at both ends. PCR products were run on a 1% agarose electrophoresis gel to confirm product size and were purified. ssRNA was transcribed from PCR products using Promega T7 RiboMAX Express kit (Cat. No. P1700) and annealed to form dsRNA. dsRNA was purified and injected into young adults at 1μg/μL concentration. Worms were dissected after 24-28hrs for imaging.

RNAi-feeding clones were obtained from the *C. elegans* RNAi library (Kamath and Ahringer, 2003), grown and sequenced for verification. Single clones were inoculated in 5 mL LB/Amp for 8 hours at 37℃ and concentrated to 1 mL. 50-100 μL was spotted on NGM plates containing 25 μg/mL Carbenicillin and 1 mM IPTG. Plates were left to dry overnight at room temperature. L4 worms were added to the plates for 24 hours and dissected for embryo imaging.

#### Drug treatment

To permeabilize the embryo’s eggshell, L4s were fed *perm-1* RNAi for 24 hours (Carvalho et al., 2011; Olson et al., 2012). Gravid adults were dissected in 10 μL Shelton’s growth media (SGM) containing Aurora A inhibitors (Figure 1, Figure S1) or PLK-1 inhibitors (Figure 4). *Ex-utero* embryos were then mounted and images as described above.

SGM was prepared by mixing the following reagents: Inulin, 1 mL of 5 mg/mL stock (Thermo, A18425.18); Polyvinylpyrrolidone powder, 50 mg (Thermo, 227545000); BME vitamins, 100 μL of 100x stock (Sigma, B6891); Chemically defined lipid concentrate, 100 μL (Gibco, 11905031); concentrated Pen-Strep, 100 μL (Sigma, P4333); Drosophila Schneider’s Medium, 9 mL (Gibco 21720024) supplemented with 35% fetal bovine serum (Gibco A3840001).

Auxin treatment was performed *ex-utero* (Figure 2) and *in utero* (Figure 3, Figure S3). For *ex-utero*, worms were first soaked in egg buffer containing 4mM auxin for 15 minutes, then dissected and mounted for imaging as described above. For *in-utero*, worms were immediately mounted in M9 containing 4mM auxin for imaging as described above.

#### Quantification and statistical analysis

Total AIR-1 fluorescent intensity was measured by drawing a region of interest (ROI) around the embryo’s perimeter (Figure 2C) or around the centrosomes (Figure 1E) and using the “analyze>measure” command in FIJI. Fluorescence intensity was obtained by subtracting off-embryo or cytoplasmic background, respectively.

Cortical PAR-3 clusters intensity in (Figure 4C) was measured by drawing an ROI at the anterior side of the embryo at the cortical plane and subtracting the off-embryo background.

The timing of events was defined relative to pronuclear meeting except where otherwise indicated. The timing of AIR-1 depletion in Figure 2E was estimated by eye because measuring fluorescent intensities *in utero* is difficult due to subtle worm movements that alter the focal plane between time-frames.

All plots were made in GraphPad Prism.

---

**Fig. S1.**
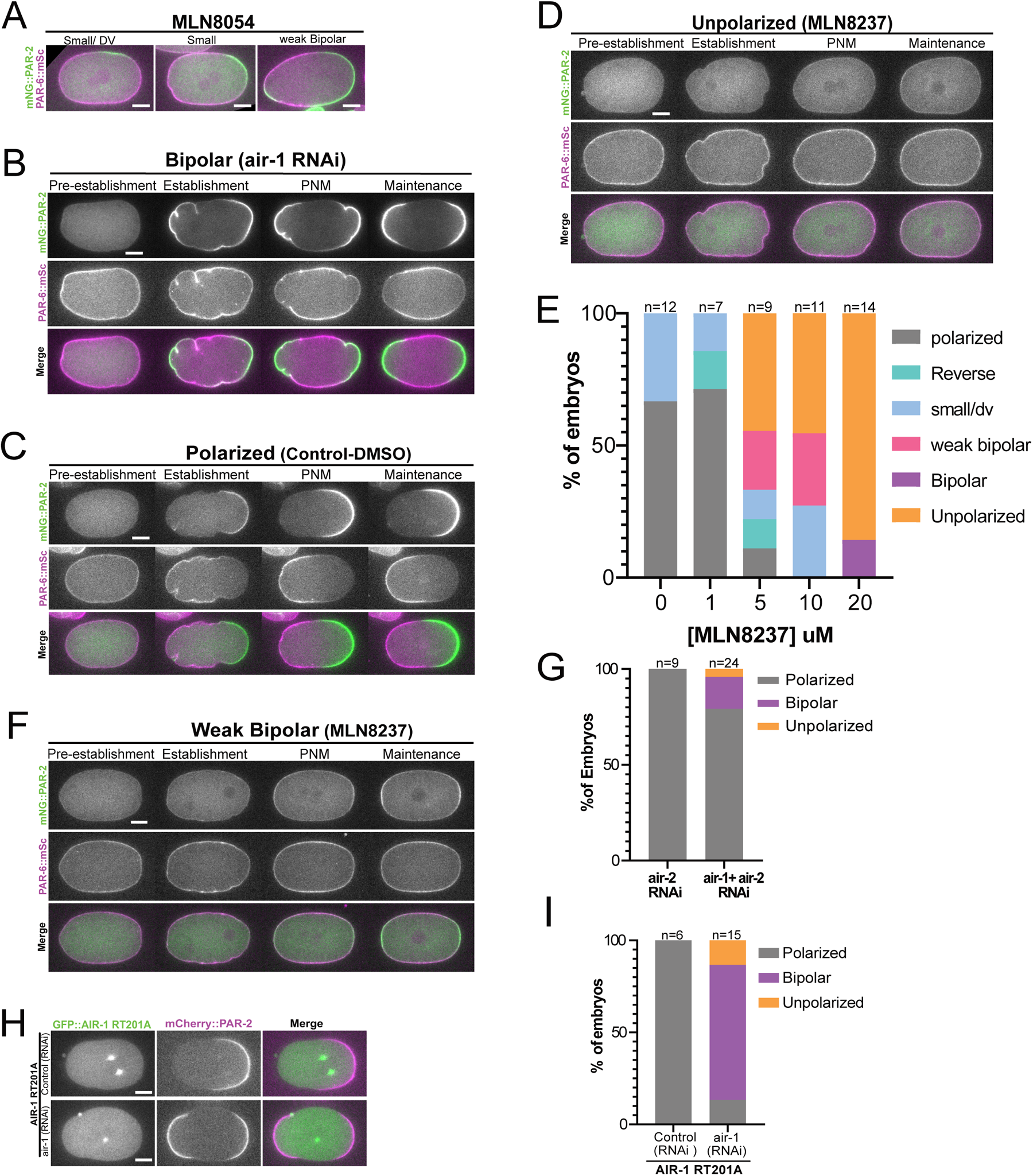
**A:** Mid-plane images from time-lapse live imaging of permeabilized mNG::PAR-2; PAR-6::mSc embryos treated with Aurora A inhibitor, MLN8054, showing different intermediate polarization defects. Scale bar represents 10μm **B**: Montage from time-lapse live imaging of *air-1 (RNAi)*; mNG::PAR-2; PAR-6::mSc embryos showing a bipolar phenotype. Scale bar represents 10μm **C-D**: Montage from time-lapse live imaging of permeabilized mNG::PAR-2; PAR-6::mSc embryos treated with 0.2% DMSO control (**C**) or Aurora A inhibitor, MLN8237 (**D**). Scale bar represents 10μm **E**: Quantification of phenotypes of embryos at pronuclear meeting (PNM) treated with Aurora A inhibitor, MLN8237 at different concentrations. **G**: Quantification of phenotypes of embryos at PNM depleted of AIR-2 alone, or AIR-1 and AIR-2 together. **F**: Montage from time-lapse live imaging of permeabilized mNG::PAR-2; PAR-6::mSc embryos treated with Aurora A inhibitor, MLN8237 showing a bipolar phenotype. Scale bar represents 10μm **H**: Mid-plane images from time-lapse live imaging of embryos expressing transgenic RNAi-resistant AIR-1::T201A::GFP; mSc::PAR-2 with or without endogenous air-1 RNAi. Scale bar represents 10μm **I**: Quantification of phenotypes scored at PNM from data in H.

**Fig. S2.**
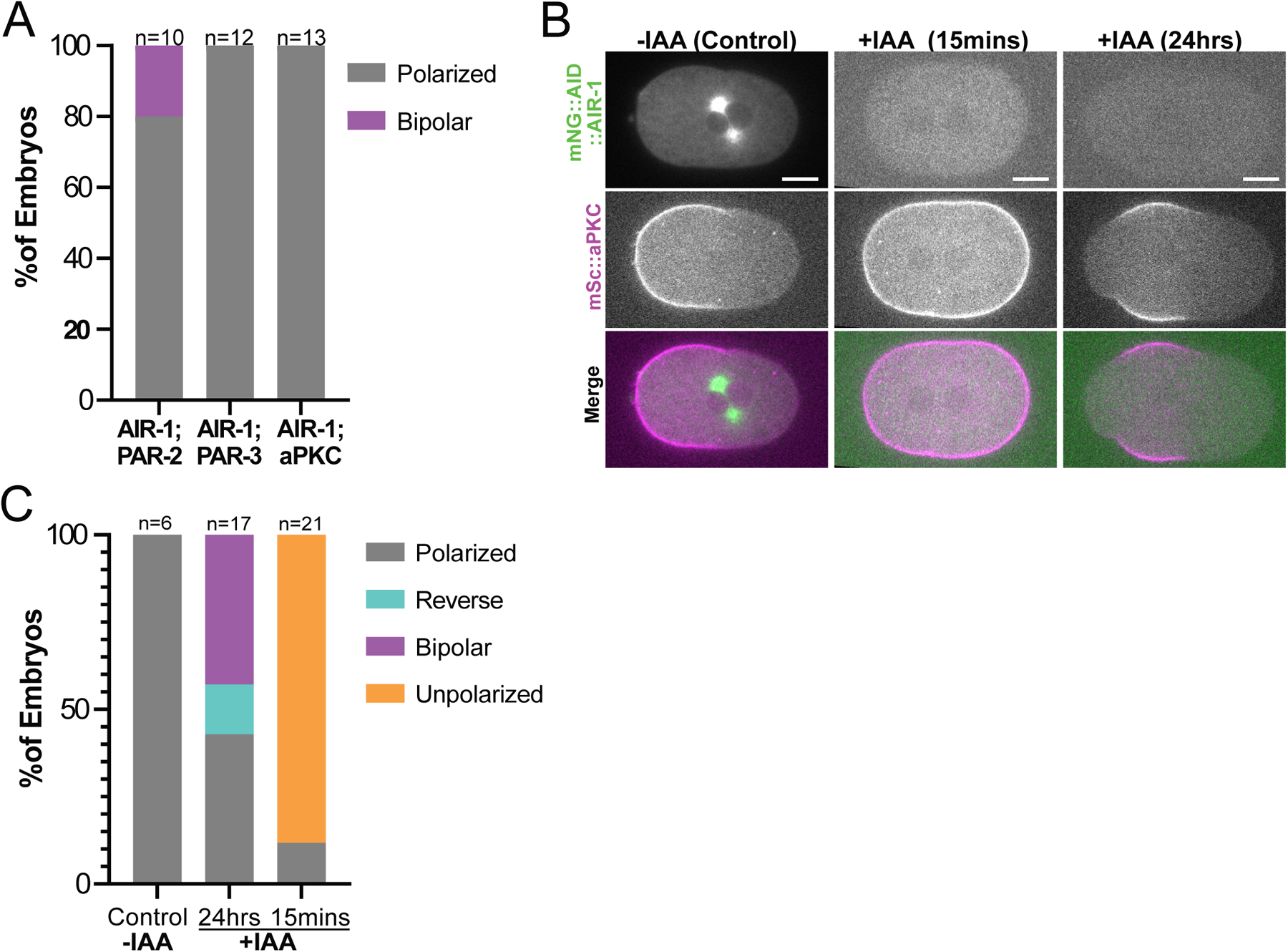
**A:** Quantification of phenotypes scored at pronuclear meeting (PNM) of strains with endogenously tagged mNG::AIR-1 in combination with mSc::PAR-2, mSc::PAR-3, or mSc::aPKC, showing different polarization phenotypes. **B**: Mid-plane images from time-lapse live imaging of mNG::AID::AIR-1; mSc::aPKC embryos treated with 1% ethanol, auxin for 15 minutes, or auxin for 24 hours. Scale bar represents 10μm **C**: Quantification of phenotype scored at PNM from panels in B.

**Fig S3.**
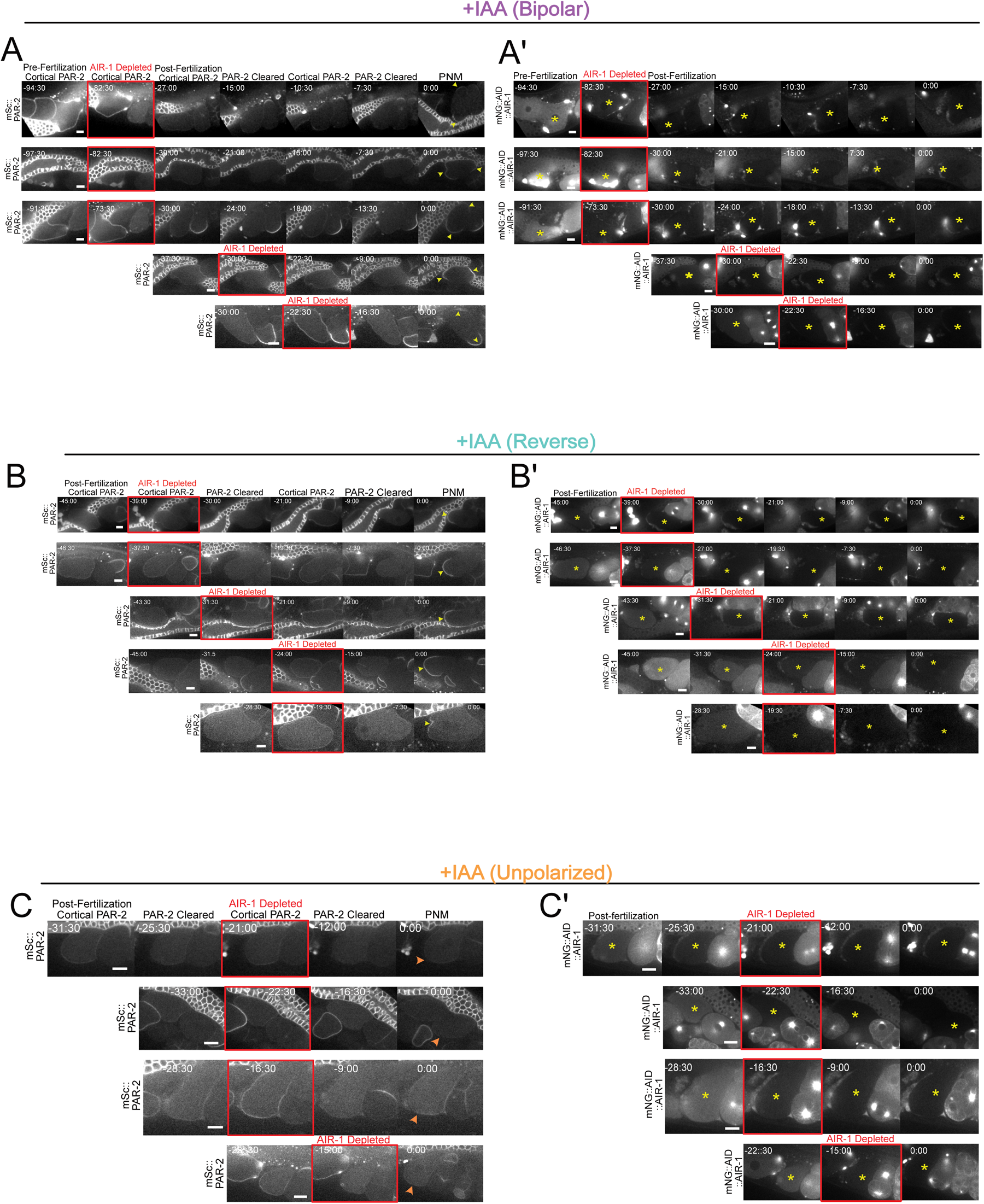
**A-C:** Additional examples of data in Figure 3 showing montages of time-lapse live imaging of embryos expressing mSc::PAR-2 (**A-C**) and mNG::AID::AIR-1 (**A’-C’**). Embryos were imaged *in utero* from worms mounted in 4mM Auxin (IAA) containing buffer. The time of AIR-1 depletion is shown in red and the resulting phenotypes after AIR-1 depletion show bipolar (**A**), reverse (**B**), and unpolarized (**C**). Yellow arrowheads indicate PAR-2 localized to the membrane, while orange arrowheads indicate embryos lacking cortical PAR-2. Yellow asterisks show the representative embryo. The timescale is in minutes relative to pronuclear meeting (PNM). Scale bar represents 10μm

